# Probe-Free Multiplexed RPA Detection Via Single-Molecule Nanopore Sensing and Deep Learning Classification

**DOI:** 10.1101/2025.04.24.650542

**Authors:** Md. Ahasan Ahamed, Muhammad Asad Ullah Khalid, Anthony J. Politza, Wahid Uz Zaman, Weihua Guan

## Abstract

Recombinase Polymerase Amplification (RPA) is a rapid, sensitive, and isothermal method for nucleic acid amplification that has gained widespread use in diagnostic applications. While well-suited for single-target assays, extending RPA to multiplex detection remains technically challenging despite the multiplexed RPA being essential for tasks such as differential diagnosis and inclusion of internal controls. Existing multiplex RPA strategies rely on proprietary probes (e.g., Exo, Fpg, Nfo), which require complex design, are susceptible to cross-reactivity, and often depend on sophisticated optical instrumentation, limiting their scalability. To address these limitations, here, we developed a probe-free multiplex RPA assay that distinguishes targets by amplicon length, using solid-state nanopore detection of single molecules and deep neural network (DNN)-based classification. Using the Monkeypox (Mpox) as a model, we designed and validated a multiplex RPA assay targeting both the Mpox gene and the human RNase P gene as an internal control. We systematically evaluated the nanopore size requirement for detecting DNA amplicons ranging from 50 to 500 base pairs and found that ∼7 nm pores provided optimal performance for amplicons between 75 and 500 bp, balancing high event rates without pore clogging. Using single-molecule translocation events as input, we trained and optimized a DNN to classify amplicons by target. The model achieved 94.2% accuracy at the single-molecule event level, which translated to 100% accuracy at the population-level target call. While our current system demonstrates duplex detection, the strategy is inherently scalable to higher multiplex levels. These findings establish a new framework for multiplex RPA that eliminates the need for complex probe design and optical detection, paving the way for robust, scalable, and accessible molecular diagnostics.

---

Recombinase polymerase amplification (RPA) is an emerging technology among other amplification techniques.^1^ RPA reactions take place at a steady temperature, typically around 37-41°C, much faster, removing the need for thermal cycling and thereby simplifying both the process and the required equipment.^2^ Other popular isothermal amplification techniques, such as Loop-Mediated Isothermal Amplification (LAMP), require complex sets of 4 to 6 primers at six binding sites.^3,4^ LAMP is unsuitable for smaller target lengths and has a higher false positive rate (over 15%).^5,6^ However, RPA leverages only two primers, which is also suitable for small-size targets,^7^ low false positive rates,^8^ and consumes less energy for its instruments than LAMP. Additionally, LAMP results in a ladder-like gel pattern due to the cauliflower-like structures of its amplicons,^9^ while RPA amplicons show a single, clear, and dense band on the agarose gel.^10^ Polymerase Chain Reaction (PCR) is another gold standard amplification method using two primers and a probe.^6^ Although PCR is highly sensitive and specific, it requires complex instrumentation and controlled temperature cycling, which also has a longer turnaround time to complete the assay. Compared to PCR, RPA tolerates 9 mismatches on primer binding sites, which gives more freedom in primer design and helps to detect slight variations in target sites.^11^ Generally, single-target detection of RPA amplicons is usually done by gel or capillary electrophoresis,^7^ lateral flow assay(LFA),^12^ and fluorescence-based detection (FBD) assisted by Exo, Fpg, Nfo, or intercalate dyes.^2,6,13^ Gel electrophoresis is limited by longer migration and analysis times, while capillary electrophoresis struggles with optimizing electrophoresis conditions, fluctuations in electroosmotic flow, and maintaining peak efficiency.^14^ Further, the LFA method lacks sensitivity and quantification, is affected by temperature and humidity, and has constrained physical limitations like cross-reactivity of biomarkers and higher background and false positives.^15,16^ The FBD method has gained attention due to its target quantification ability, sensitivity and specificity, lower detection limit, and multiplexing ability.^17^

Multiplexing of target amplicons is highly desirable, as it enables the simultaneous detection of multiple pathogens, such as respiratory infections (SARS-CoV-2, respiratory syncytial virus, influenza),^18^ flavivirus infections (Zika virus, dengue virus),^19^ and skin infections (monkeypox, chickenpox, smallpox),^20^ while conserving reagent volumes, reducing assay time, and lowering overall diagnostic costs.^21^ While the use of intercalating fluorophores such as SYBR Green and EVA Green can simplify FBD,^22^ these dyes are inherently prone to generating nonspecific background signals,^2^ compromising assay specificity. To overcome this limitation, probe-based strategies employing sequence-specific probes (e.g., Exo, Fpg, and Nfo probes) are utilized for multiplexed FBD reactions.^23^ However, this approach introduces additional challenges, including increased susceptibility to cross-reactivity and a reliance on sophisticated optical instrumentation, which restricts scalability.^2,24–26^ Moreover, conventional TaqMan probes are incompatible with RPA because the 5’-3’ exonuclease activity of Taq polymerase degrades displaced DNA strands during strand displacement.^10^ An additional obstacle lies in the limited multiplexing capacity of optical systems, which is constrained by the spectral overlap of fluorophores, difficulties in selecting appropriate wavelength LEDs, and the necessity for multiple optical filters.^27–29^ These factors collectively impose significant barriers to scalability. Recently, electronic-based detection methods such as electrochemical sensors,^30^ solid-state nanopore (SSN) sensors,^7,31,32^ and field-effect transistors^33^ have shown great promise for molecular diagnostics due to their potential for integration and miniaturization. Among them, nanopore sensing offers unique advantages for single-molecule detection, including a probe-free approach and the ability for digital sizing and counting.^5^ SSN sensors, in particular, have been adapted for multiplex detection through strategies like probe-encoding^34^ and length-encoding^35^. Moreover, advancements in post-acquisition data analysis using machine learning (ML) have significantly enhanced target detection accuracy by filtering out background noise and mitigating pore-to-pore signal variation.^5,36–38^ In our previous study, we developed an RPA-CRISPR strategy to identify RPA amplicons by utilizing 3.5 kbp ssDNA reporters, where specific detection relied on CRISPR-mediated cleavage activity.^7,39^ However, this indirect detection method introduced an additional CRISPR step and increased the overall turnaround time. We initially integrated CRISPR because, with ∼10–12 nm nanopores, we could not confidently and reproducibly detect the ∼117 bp Mpox amplicon. These limitations highlight the critical need for direct detection of short-sized RPA amplicons using SSN sensors to advance rapid, robust, and multiplexed molecular diagnostics. Therefore, there is an unmet challenge to develop an SSN-based method capable of reliably detecting multiplex RPA amplicons with high sensitivity and reproducibility.

In this work, we developed a deep neural network (DNN)-assisted method to classify multiplex RPA products using a probe-free, length-encoded electronic readout from an SSN sensor. We validated a multiplex RPA assay with Mpox as a model, detecting both the Mpox target gene and the human RNase P gene as an internal control. We then optimized nanopore size for efficient DNA amplicon detection, balancing sensitivity and minimizing pore clogging. Finally, we leveraged single-molecule translocation events to train and refine a DNN capable of classifying amplicons based on individual events. The model demonstrated strong performance at the individual event level, leading to accurate identification at the population scale. While our current platform demonstrates duplex detection, it is readily scalable, establishing a new framework for multiplex RPA that eliminates complex probe designs and optical systems and advances robust, scalable molecular diagnostics.

## RESULTS

### Principles of DNN-based multiplex RPA classification

To address the need to resolve two populations of RPA amplicons by length using nanopore technology, we conducted a study in which a 110 bp Mpox target was selected as the model and the human RNase P gene as an internal control, both chosen to balance assay design and nanopore detection considerations. In our experiment, we employed an RPA amplification reaction where DNA of different lengths, Target 1 (T_1_) and Target 2 (T_2_), are shown in **Figure 1a**. Two targets are amplified in the one-pot buffer and produce a lot of amplicons. Then, the generated amplicons are passed through the nanopore. The primary challenge lies in the detection capabilities of nanopores, as previous works indicate a minimum detection limit of around 200 bp using more than 10 nm bare glass nanopores.^40^ However, with specific surface modifications such as introducing a gel structure inside the nanopore, the resolution can be improved to around 50 bp.^41,42^ To solve the sensitivity and resolving capability issue, we chose the length encoding of L_1_ and L_2_ of both targets and utilized ∼7 nm diameter glass nanopores for measurement. The resulting ionic current-time traces (*I-t*) are recorded and analyzed for targets T_1_ and T_2_. In **Figure 1b**, our findings statistically demonstrate that the nanopore struggles to distinguish both targets with distinct distributions, as the data show overlapping current peaks between 25 and 50 pA and dwell times of less than 500 µs. For clearer visualization, the contour plot is shown in Supplementary **Figure S1**, revealing a significant overlap in the 15–25 pA range.

**Figure 1.**
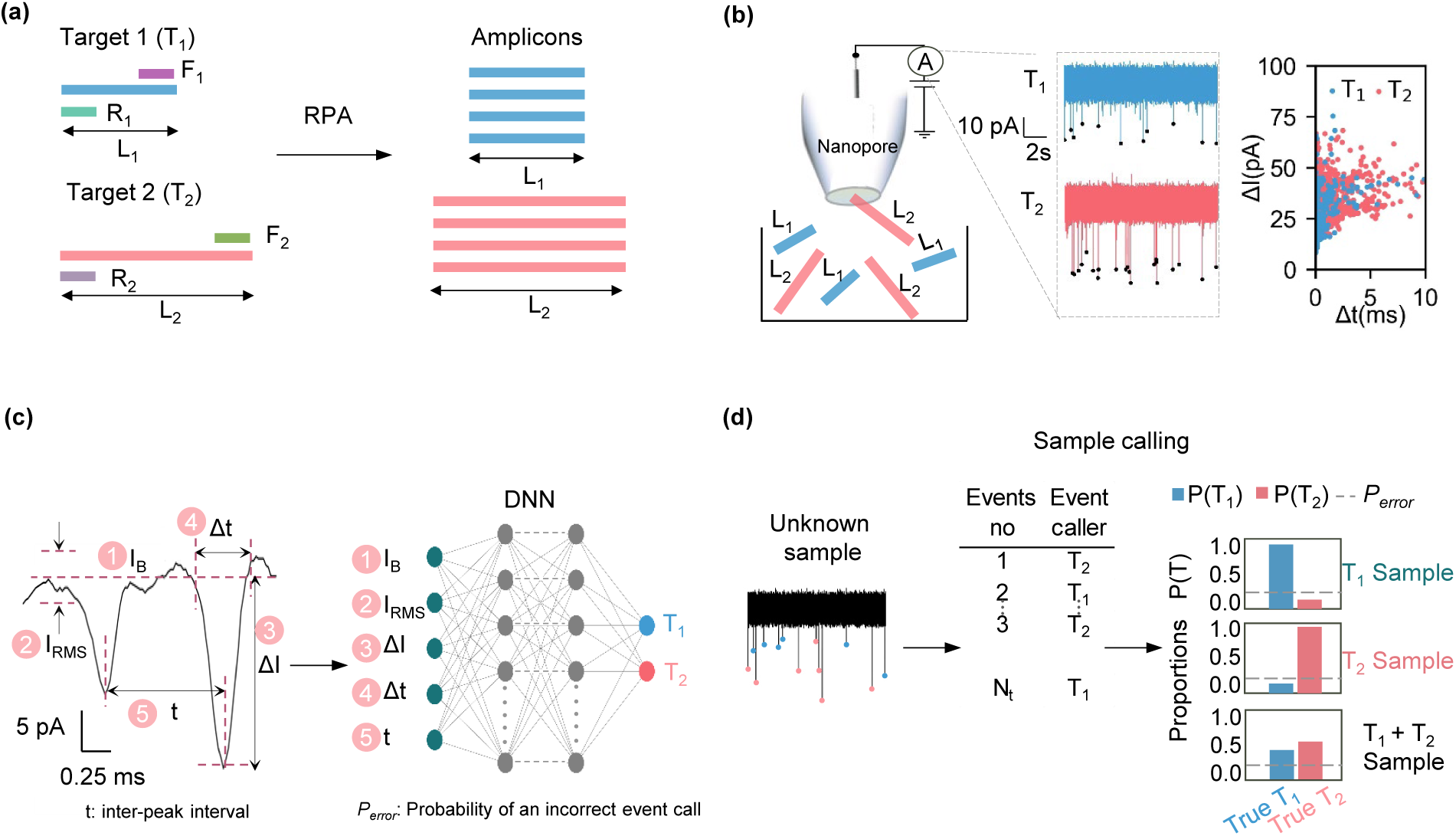
Workflow of Multiplex RPA Amplicon Sensing. **a)** RPA reaction amplifies two length-encoded targets (T_1_ and T_2_), with distinct amplicon lengths for differentiation, enabling simultaneous detection. **b)** Nanopore sensing captures event waveforms corresponding to RPA amplicons, generating signals characterized by multiple parameters. **c)** Key waveform features extracted include baseline current (I_B_), RMS current (I_RMS_), blockage current (ΔI), dwell time (Δt), and inter-event distance (t). These features train a Deep Neural Network (DNN) model that outputs a classification error probability. **d)** For unknown samples, the DNN model classifies each event individually as T_1_ or T_2_, determining the Proportion P(T) of T_1_ or T_2_ or T_1+2_ target presence.

Given these challenges, we incorporated deep neural network (DNN) into our analysis to improve the resolution and accuracy of detecting DNA populations. DNN algorithms were trained to identify subtle differences in the ionic current traces that may not be easily distinguishable through traditional statistical analysis methods such as event charge deficit (ECD) distribution, blockage and dwell time counts distributions, interarrival time distributions, etc. This approach allowed us to better differentiate between closely sized DNA molecules by learning from a large dataset of nanopore signals. In contrast, our DNN was trained on five extracted features from the ionic current traces: baseline current (I_B_), RMS current (I_RMS_), signal blockage (ΔI), dwell time (Δt), and inter-peak interval (t). These features were derived from the event waveform analysis of two known classes, T_1_ and T_2_. In **Figure 1c**, we illustrate the five features used as input to the DNN model. Specifically, I_B_ and I_RMS_ account for pore-to-pore variability, while ΔI and Δt are influenced by the size and shape of the DNA amplicons. The fifth feature, t (time between two consecutive peaks), helps characterize event frequency and signal dynamics, supporting discrimination between no-template control (NTC) and positive control (PC) samples. Supplementary **Figure S2** shows the extracted waveform of each signal, where the first four features (I_B_, I_RMS_, ΔI, and Δt) are depicted in detail. This feature-based learning allowed the DNN to accurately classify complex signals and significantly improved the reliability of RPA amplicons detection using our nanopore-based assay. After training, we tested the model on test samples and evaluated the probability of incorrect event classification (*P_error_*). Finally, the trained model was applied to completely unknown samples (those not included in the initial dataset) for classification, as shown in **Figure 1d**. The model operates at the single molecular level, analyzing each event individually. For every detected event (1, 2, 3, …, N), it predicts whether the signal corresponds to T_1_ or T_2_ and calculates the proportion P(T), the likelihood of that event belonging to either class, using a decision threshold derived from the *P_error_* observed during testing. Based on the aggregated P(T) values across events, we assigned a final population label to each sample: T_1_, T_2_, or T_1_ + T_2_. This workflow enables robust classification of unknown samples and presents a promising advancement for the accurate detection of multiplex RPA amplicons, supporting broader application to infectious disease diagnostics at the point of care (POC).

### Length-coded RPA multiplex assay

To develop an RPA multiplex assay, it is critical to accurately distinguish between multiple target lengths, which are target T_1_ and T_2_ of length L_1_ and L_2,_ respectively. It involves targeting the F3L gene fragments of the 2022 USA MPXV strain, specifically the 46,344 bp to 46,453 bp region Human RPP30 (RNase P) gene, specifically the 100 bp to 309 bp region in **Figure 2a**. The primer sets (F_1_ and R_1_) for Mpox were redesigned from our last work by slightly modifying the target length, which was initially 117 bp.^7,39^ The detailed sequences are shown in Supplementary **Table S1**. Rnase P primers (F_2_ and R_2_) are selected from the combination of three forward primers and four reverse primers using PCR reaction, shown in Supplementary **Figure S3a**. The RFU plot in Supplementary **Figure S3b** was used to screen the RNase P sequence, where two primer sets (F_2_R_2_ and F_d_R_2_) exhibited high-endpoint RFU signals and identical Cq values. We analyzed the theoretical ΔG (kcal/mol) values to further evaluate potential cross-reactivity, with a threshold of -6 kcal/mol indicating cross-reactivity^43^. Since both primer sets showed similar ΔG values, we selected F_2_R_2_, which had the highest end-point RFU signal. To ensure the cross-reactivity of primers of Mpox and RNase P, we run additional gel electrophoresis. In Supplementary **Figure S4**, the gel lane 1-4 indicates the primer’s combination of F_1_R_1_, F_1_R_2_, F_2_R_1_, and F_2_R_2_. Faint light bands were observed similar to NTC (gel lane 5), confirming no possible cross-reactivity between primers. Building on this, in **Figure 2b**, we selected primers F_1_, F_2,_ R_1,_ and R_2_ for RPA multiplex amplification reaction of two different lengths, L_1_ and L_2_, to demonstrate the feasibility of multiplexing these targets. The amplified products were then analyzed using gel electrophoresis to confirm successful and specific amplification.

**Figure 2.**
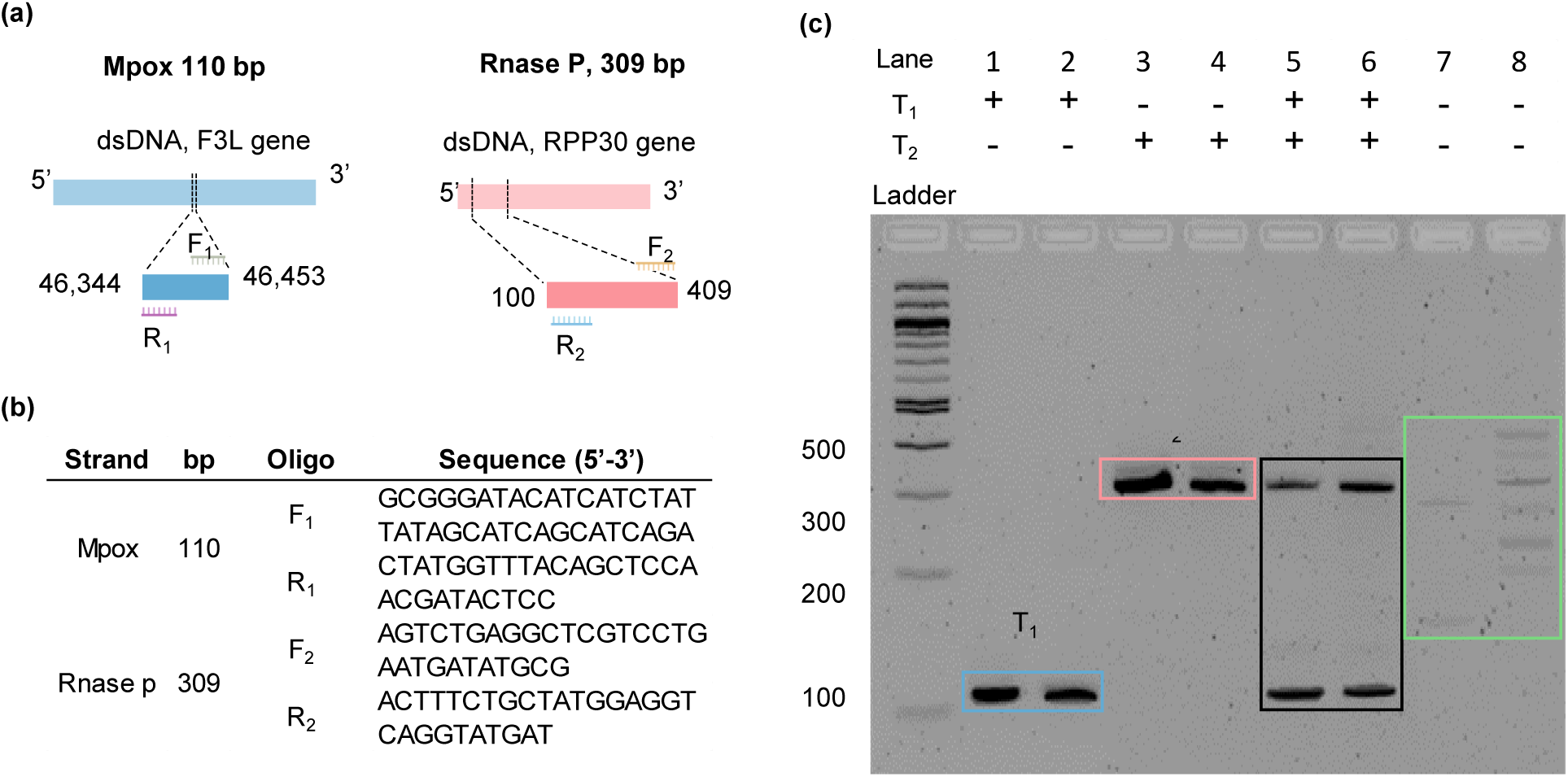
Validation of the multiplex RPA assay. (a) The selected genes for Monkeypox (Mpox) and human RNase P DNA serve as process controls. The multiplex assay includes forward primers (F_1_, F_2_) and reverse primers (R_1_, R_2_). (b) Oligonucleotide sequences of both primer sets. (c) Gel electrophoresis validation of the multiplex assay, where the first two lanes correspond to T_1_, the next two to T_2_, followed by two lanes representing the multiplex assay (T_1_ + T_2_), and the last two lanes containing the NTC samples.

To investigate the outcome of the multiplex assay, the gel electrophoresis results (**Figure 2c**) confirm that our primers successfully amplified target T_1_ and T_2_ with distinct bands for each target. The mixed sample (T_1_ + T_2_ ) shows bands corresponding to both targets, indicating that the multiplex assay can amplify multiple targets simultaneously. However, faint bands are also lightly observed in lane 6 and lanes 7 and 8 of NTC, which means that there are possible artifacts in the RPA assay. Given the limitations of artifacts in the RPA assay, these results validate the feasibility of the RPA multiplex assay for detecting multiple targets, which is essential for advancing nanopore-based detection. The detailed recipe for singleplex and multiplex reactions is provided in Supplementary **Table S2** and Supplementary **Table S3,** respectively. Based on these results, we are confident in proceeding with the nanopore experiment for the length-encoded target, as the amplification of both targets was successful in the multiplex assay.

### Sensitivity of nanopore sensor

To reliably detect multiplexed RPA products and distinguish them from background signals, it is essential to accurately capture translocation events across the full expected amplicon size range (80–500 bp).^44^ This analysis also informs the selection of suitable nanopore sizes for effective detection of short-size DNA. The success of multiplexing reactions hinges on the events picked up by the nanopore sensor for 100 bp and 300 bp DNA fragments. Previously, Li et al. demonstrated a bare glass nonopore size resolution of about 200 bp but did not explore the sub-10 nm range or sizes below 200 bp.^40^ We analyzed the impact of nanopore size on detecting various DNA fragment sizes from 50 bp to 500 bp. To evaluate the suitability of different nanopore sizes for detecting RPA amplicons, we examined the event frequency with varying nanopore diameters from 3.4 nm to 12.4 nm with the DNA fragments concentration of 1 nM, which encompasses the possible lengths of RPA amplicons. The nanopore diameter (*d*) is calculated using a simplified conical pore model:^45^ 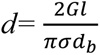, where *G* is the measured ionic conductance (nS), *σ* is the electrolyte conductivity (S/m), *d_b_* is the base diameter of the nanopore (µm), and *l* is the nanopore length (µm). The detailed formulae are derived in Supplementary **Section 1**. The current–time (*I–t*) traces captured by nanopore sensors reveal three distinct phases in the sensing process: a clogging region, a less sensitive region, and a sensing region, as illustrated in **Figure 3a**. The clogging chance increases if the radius of Gyration, *R* of DNA size, is much bigger than the nanopore diameter, which *R^2^* = *L*×*P*/3 indicates, *L*= DNA Contour length (*N*×0.34 nm), *N*= size (bp), *P*= Persistence length^4,46^. Translocation typically occurs through a stochastic threading process.^47^ In Supplementary **Figure S5**, physical phenomena that arise during sensing as both nanopore diameter and DNA fragment size vary. For longer DNA fragments such as 300 bp and 500 bp, with corresponding *R*-values of 41.2 nm and 53.2 nm, respectively, the likelihood of clogging increases significantly when small nanopores (<5.2 nm) are used. Conversely, for smaller fragments like 75 bp and 100 bp, the use of large pores (>8.6 nm) results in high translocation velocities, causing many events to go undetected and thereby reducing sensor sensitivity. Notably, **Figure 3b** highlights that nanopores around ∼7 nm showed a critical balance. This size range enables effective detection of amplicons between 75 and 500 bp, supporting high event rates via threading without significant pore clogging. In the heat map, the green color shows the area of detection, and the red color shows the area of undetected events and clogged events. The minimum possible size of RPA amplicons is approximately 80 bp.^44,48^ To clarify and validate whether our sensor could accurately detect such small amplicons rather than background noise, we conducted an additional experiment using 75 bp DNA fragments. Measurements were performed in 1 M KCl with a 6.8 nm diameter nanopore and a final solution volume of 200 µL to enhance conductivity. The resulting event frequency is shown in Supplementary **Figure S6**. Initially, as the nanopore diameter increases, event frequency rises because DNA molecules pass through more easily. However, further increases beyond a certain threshold reduce event frequency due to weaker and less specific interactions between DNA and the nanopore, compromising the nanopore’s ability to capture and translocate DNA effectively. With these results, we gain a clear understanding of ∼7 nm nanopore diameter required for effective signal detection across multiple nanopores, which is essential for acquiring the large datasets needed for DNN training and testing.

**Figure 3.**
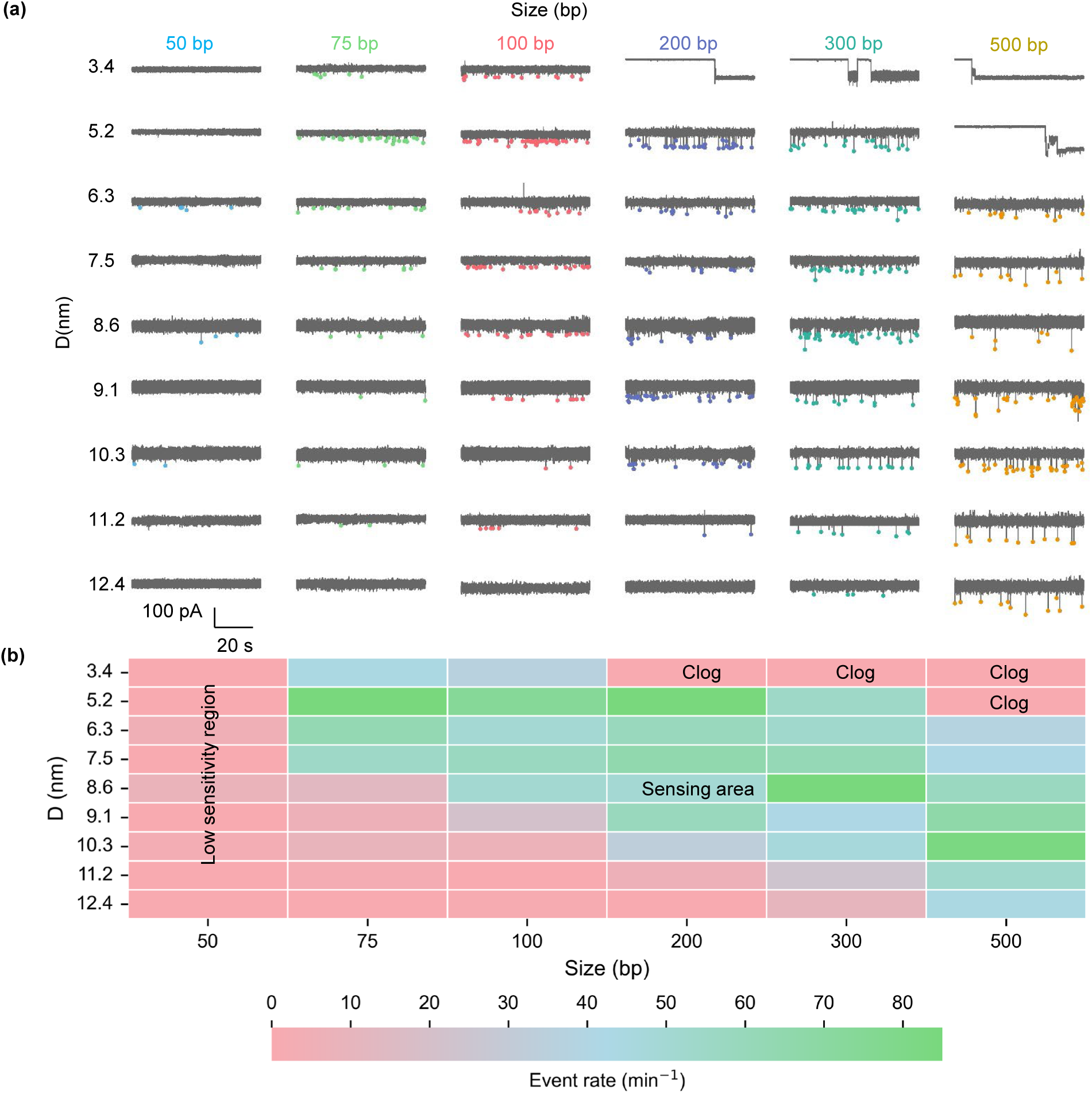
Sensitivity of the Nanopore Sensor. (a) Current trace (I–t) for 1nM model DNA fragments ranging from 50 to 500 bp, corresponding to nanopore diameters between 3.4 nm and 12.4 nm. (b) Sensitivity analysis illustrating the sensitive, less sensitive, and clogging zones, with a color map representing the event rate (min⁻¹).

### Training of Deep Neural Network (DNN) model

To precisely distinguish multiplexed RPA targets from SSN sensor data, DNN is essential for accurately interpreting overlapping signals and overcoming the statistical challenges of differentiating targets T_1_ and T_2_. As shown in **Figure 2c**, the NTC data contains some background signals generated by RPA reactions (lanes 7 and 8 of gel image); however, a 20× dilution under high-salt conditions minimizes these signals while maintaining an optimal event rate, reducing the number of artifacts, as shown in Supplementary **Figure S7**. We avoided a 10× dilution to prevent clogging, whereas a 40× dilution led to an excessively slow event rate. The inter-peak interval, t (i.e., time between two consecutive NTC events), is very large. Based on this insight, we defined the features of the DNN model described in **Figure 1c**, where the final feature helps distinguish NTC signals from other PC signals. To collect data, we measured events for 5 minutes from each target (T_1_ and T_2_) across 15 nanopores (totaling 30 samples) using a threshold (μ + nσ), where μ and σ represent the mean baseline current and current drop from the baseline, respectively. We chose n = 3 to 10, depending on pore noise, and maintained a unique value for all targets within the same pore. Because clogging occurred during sensing, we omitted clogging portions from the analysis, so not all samples had a full 5 minutes of effective data. We then used 30 samples from T_1_ and T_2_ for DNN model training. Initially, we evaluated nine ML models using 301 test data collected from two nanopores (Supplementary **Figure S8**), where 1500 data points were used for training from 8 nanopores. Based on the nature and complexity of the data, we found that the DNN model consistently outperformed other ML models, including Random Forest, Histogram-Based Gradient Boosting (HGB), Gradient Boosting, Decision Tree, K-Nearest Neighbors (KNN), Support Vector Machine (SVM), Logistic Regression, and Naïve Bayes. While it is true that all models can achieve high accuracy depending on training quality and parameter tuning, the DNN model demonstrated superior performance with the existing dataset and provided better generalization. Therefore, we selected the DNN model as the primary model for our study. **Figure 4a** shows the architecture of our DNN, which has an input layer and five hidden layers with 512, 256, 128, 64, and 32 nodes, respectively, which are fully connected, and an output layer. We used ReLU activation for each of the five hidden layers, softmax to convert raw output scores (logits) into probability distributions summing to 1, thereby assigning confidence scores to each class for binary classification, and applied a cross-entropy loss function. **Figure 4b** illustrates the overall process flow for data/model preparation, model training, and testing, covering everything from data loading and tensor creation to final classification. The workflow begins with data preparation, where the dataset is imported, split into training, validation, and test sets, and normalized to ensure consistent feature scaling, as illustrated in Supplementary **Figure S9**. In the data loading and processing stage, the prepared data is converted into PyTorch tensors to ensure compatibility with the framework. It is then organized into batches using a DataLoader, which optimizes memory usage and enables parallelized computation during training. During training, the model iterates through several epochs; each epoch involves a training step, forward passes generate predictions, cross-entropy loss is computed, gradients are derived via backpropagation, and the optimizer updates model weights, followed by a validation step assessing performance metrics (accuracy) and dynamically adjusting the learning rate if necessary. After training, the final model is evaluated on a designated test set to measure generalization. Moreover, we iteratively refine the neural network architecture and fine-tune its settings, repeating the entire process until the model stops improving. While the current task is a binary classification problem with two target classes, the model is designed as a multiclass classifier.

**Figure 4.**
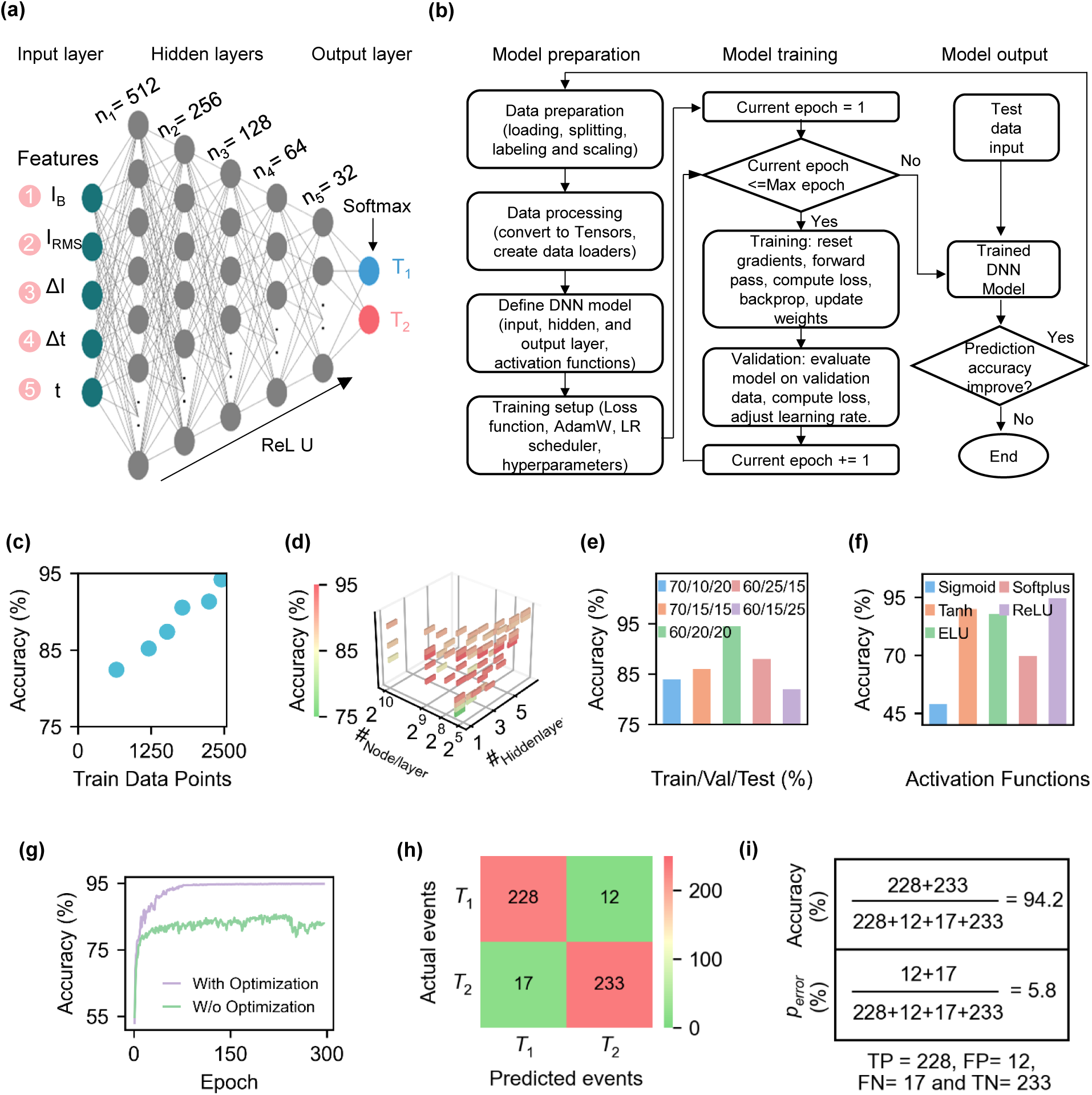
Deep Neural Network Model. a) The schematic diagram of the DNN model consists of input features, hidden layers, and node distribution, utilizing ReLU and Softmax activation functions. b) End-to-end workflow of DNN model development: model preparation (left), model training (middle), and model output (right), progressing left to right. c) A total of 2,450 data points were used for training. d) Various hidden layer and node configurations were tested, with the best-performing setup comprising five layers with 512, 256, 128, 64, and 32 nodes per layer, achieving an accuracy of 94.2%. e) Data was split into 60% training, 20% validation, and 20% testing, optimizing model performance with the selected five-layer architecture. f) Different activation functions were compared within the hidden layers to assess their impact on accuracy. g) Model performance was evaluated with and without the Adaptive Moment Estimation (ADAM) optimizer. h) Testing was performed on 490 data points (20% of the dataset). i) The final accuracy was 94.2%, with an error rate of 5.8% for incorrect event classification.

To understand how many data points are needed to achieve clinically relevant accuracy, **Figure 4c** explores the required number of data points where the 30 samples contained 2450 data points. Current traces of all 30 samples are shown in Supplementary **Figure S10**, which can give the idea pore-to-pore data variations for picking up signals of both targets. We evaluated various hidden-layer and node combinations (**Figure 4d**) and determined that the configuration shown in **Figure 4a**, consisting of five hidden layers with 512, 256, 128, 64, and 32 nodes, achieved the highest accuracy. In Supplementary **Table S5**, the accuracy for each combination of hidden-layer and node is shown. **Figure 4e** shows a bar plot examining how different data splits affect model accuracy, revealing that a 60%/20%/20% (train/validation/test) split yields the best performance. To avoid the vanishing gradient problem, which is particularly important since nanopore data can have both small and large values, we evaluated several activation functions (sigmoid, softplus, tanh, ELU, and ReLU) in **Figure 4f**, finding that ReLU outperforms the others by preventing vanishing gradients during backpropagation. Next, to reduce data oscillation and achieve faster convergence, we employed the AdamW optimizer, an enhanced version of Adaptive Moment Estimation (Adam) with weight decay for regularization and a learning rate scheduler (ReduceLROnPlateau) that lowers the learning rate when validation performance plateaus (**Figure 4g**). This combination boosted accuracy by roughly ∼10% compared to training without AdamW, as the adaptive learning rates helped accelerate convergence and stabilize training. Finally, to assess the model’s performance, we tested 492 events of T_1_ (TP) and T_2_ (TN) (**Figure 4h**), obtaining 12 false positives from T_1_ and 17 false negatives from T_2_, corresponding to an accuracy of 94.2% and an error probability (P_error_) of 5.8% (**Figure 4i**). Although there is always room for improving accuracy through architecture modifications and hyperparameter tuning (as described in **Figure 4b**), these results demonstrate that we can now confidently apply this model for unknown sample classification.

### Testing of DNN model for unknown samples

To evaluate the DNN model performance for unknown sample detection, we applied our trained DNN model and also explored unknown (unseen) data analysis. We conducted experiments using 19 additional nanopores and 35 new samples that were not used during training; with an inherent 5.8% error rate (**Figure 4i**), the model predicted each event individually at single molecular level (**Figure 1d**). We defined the total number of events as *N_t_* with *N_1_* and *N_2_* representing the number of events classified as T_1_ and T_2_, respectively. The proportion of true classification P(T) reflects the corrected assignment of each class. Assuming the probability of correct classification is (1 − *P_error_*), the expected number of correctly classified events for each class is *N_1_*(1 − *P_error_*) for T_1_ and *N_2_*(1 − *P_error_*) for T_2_. Misclassifications are represented by *N_1_P_error_* + *N_2_P_error_*. In a pure T_1_ sample, almost all events are classified as T_1_ and very few as T_2_; the Proportion of T_1_ approaches one, and that of T_2_ approaches zero. Using this framework, we calculated the corrected proportions for each class. For instance, the corrected proportion of T_1_ sample: 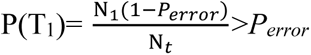, 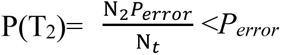. Similarly, for T_2_ sample: 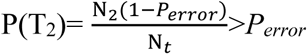, 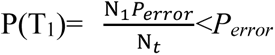 and mixed T_1_+ T_2_ sample: 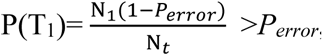, 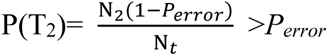. A sample is considered T_1_ if the proportion of P(T_1_) is greater than *P_error_*, and P(T_2_) is less than *P_error_*. Conversely, a T_2_ sample shows the opposite pattern. A mixture of T_1_ and T_2_ is detected when both corrected proportions exceed the error rate. These details are derived in **Supplementary Section 2**. Based on this framework, in **Figure 5a**, we evaluate the proportion distribution of blinded test samples (they are not included in the initial training, validation, and testing, consisting of 11 T_1_, 9 T_2_, and 15 T_1_+T_2_ samples. Using predefined probability thresholds *P_error_* and 1-*P_error_* samples were classified into three groups: T_1_, T_2_, and T_1_+T_2_. We found that the Proportion P(T) showed population-level classification achieved an overall accuracy of 100%. The output of P(T) is shown in Supplementary **Table S6**.

**Figure 5.**
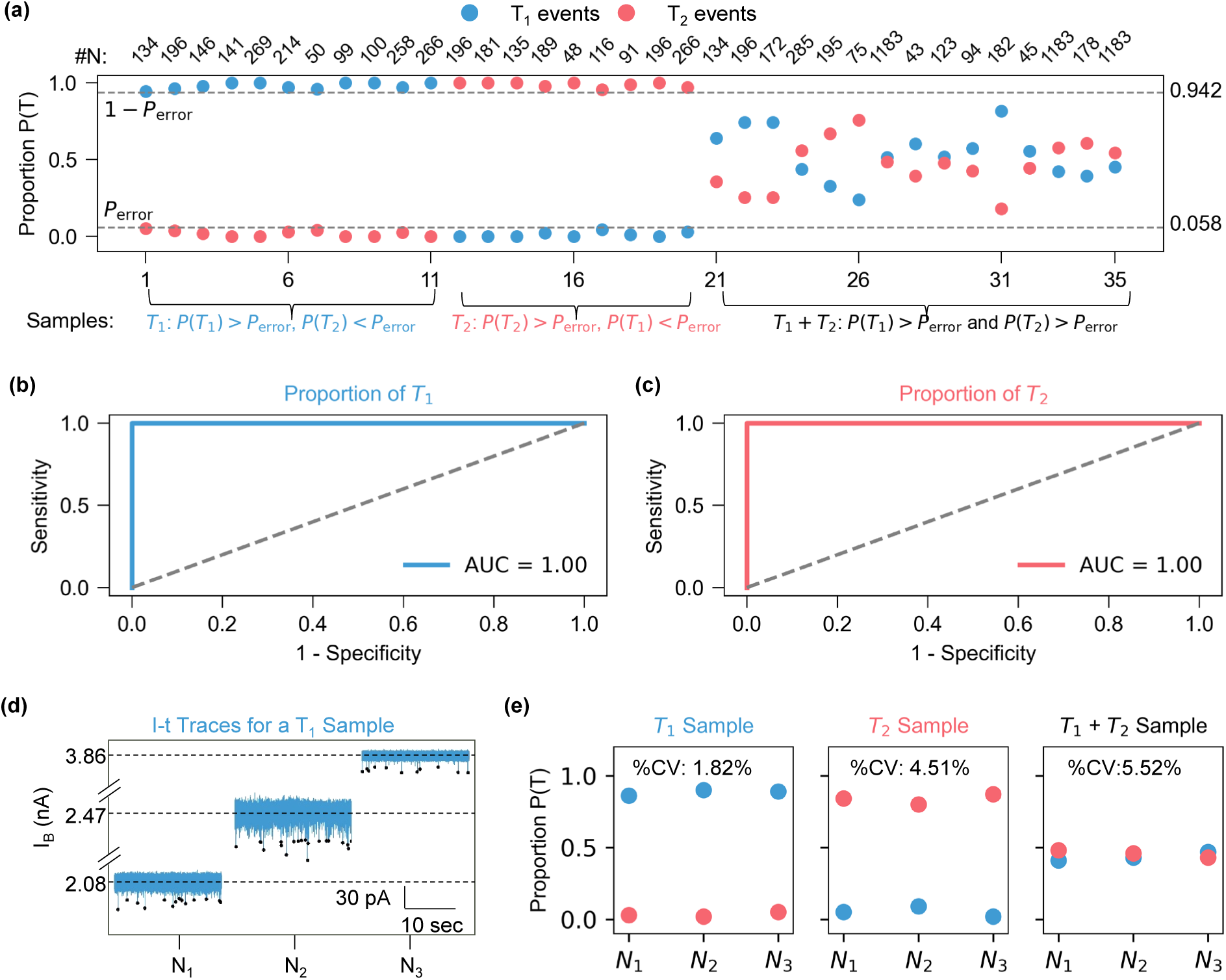
DNN Model Performance for Unknown Sample Classification. a) Panel shows the event-level proportion distribution of 35 blinded test samples classifications with proportion P(T), with each sample classified based on the proportion of T_1_ (blue) and T_2_ (red) events. The thresholds P_error_ and 1-P_error_ define the classification zones: T_1_, T_2_, and T_1_+T_2_ based on population level classification. b) ROC-AUC analysis for T_1_ classification with AUC= 1.00. c) ROC-AUC analysis for T_2_ classification, also showing AUC = 1.00. d) Representative ionic current (I–t) traces for a T_1_ sample demonstrate stable blockade levels across molecular translocation events, with minimal signal variation observed across nanopore sizes of 5.45 nm, 6.34 nm, and 7.25 nm. e) Triplicate classification results for T_1_, T_2_, and T_1_+T_2_ samples using three independent nanopores (N₁, N₂, N₃). Coefficient variation (%CV) values confirm high reproducibility and nanopore-size-independent classification accuracy.

To evaluate diagnostic performance, we computed receiver operating characteristic (ROC) curves and area under the curve (AUC) values. In **Figure 5b**, the ROC curve for T_1_ classification was generated by treating 26 T_1_ and T_1_+T_2_ samples as positive and 9 T_2_ samples as negative. The resulting AUC of 1.00 indicates perfect classification performance for T_1_. Similarly, **Figure 5c** shows the ROC curve for T_2_, where 24 T_2_ and T_1_+T_2_ samples as positive and 11 T_1_ samples as negative. This also yielded an AUC of 1.00, demonstrating excellent model accuracy for detecting T_2_, which means the callisifaction is accurate in the population lavel. To assess the influence of nanopore dimensions, **Figure 5d** presents representative ionic current (I–t) traces from a T_1_ sample across three nanopores with diameters of 5.45 nm (N₁), 6.34 nm (N₂), and 7.25 nm (N₃). The traces demonstrate clear and stable blockade patterns, with RMS noise levels of 8.4 pA, 11.2 pA, and 5.3 pA, respectively. Finally, **Figure 5e** summarizes triplicate classification results for T_1_, T_2_, and T_1_+T_2_ samples measured across the three nanopores. The coefficient of variation (%CV) values— 1.82%, 4.51%, and 5.52%—confirm high reproducibility and classification accuracy independent of nanopore size. These findings establish a new framework for multiplex RPA, paving the way for robust, scalable, and accessible molecular diagnostics that can be extended to detect multiple co-infections and enable precise molecular classification.

## DISCUSSION

RPA is an emerging isothermal amplification method operating at 37–41°C with fewer primers. Unlike LAMP and other isothermal methods, RPA can be used for short-sized targets and produce fewer false positives. Compared to PCR, which needs thermal cycling and complex equipment, RPA is simpler and more suitable for POC use. Conventional singleplex RPA amplicon detection methods, such as gel or capillary electrophoresis, lateral flow assays (LFA), and fluorescence-based detection (FBD), each face limitations in speed, sensitivity, quantification, or stability, though FBD stands out for its multiplexing capability and analytical performance. Multiplex FBD relies on expensive probes and is limited by spectral overlap, LED selection challenges, and the need for multiple optical filters. To overcome these, this study introduces a novel, probe-free, ML-aided detection strategy using SSN sensors. A multiplex RPA assay was developed for detecting model target Mpox and internal control human RNase P genes, with the amplicons length encoding. A DNN-based classifier was trained to recognize and classify targets from the events of SSN signals, enabling accurate identification of single and multiplex targets (**Figure 1**). This method advances rapid, multiplexed, and miniaturized POC diagnostics by eliminating the need for complex probes or optical systems.

Assay development is a key step in multiplexing. It requires careful primer design to prevent cross-contamination, along with optimization of primer and target concentrations to ensure efficient competition within the bulk reaction mix. Nanopore sizing resolution presents a major bottleneck for scalable multiplex detection, as conventional size-based sensing struggles to distinguish amplicons with less than 100 bp difference. To address this, we adopted a length-encoding strategy, ensuring at least a 100 bp size gap between targets.^41^ We developed a duplex RPA assay using Mpox and RNase P. RNase P can also be substituted with any co-infection target. As shown in **Figure 2c** (lanes 5 and 6 of gel image), we successfully developed a multiplex RPA reaction compatible with nanopore sensing.

Next, we focused on determining the optimal performance of the nanopore sensor to detect the full RPA amplicon size range. Detecting small RPA amplicons is often challenging due to high drift forces, which can either accelerate translocation or delay detection. To enhance sensitivity, we reduced the nanopore diameter to below 10 nm. Sub-10 nm glass nanopores, particularly those around ∼7 nm, exhibited high event frequency without clogging. This led us to identify an optimal diameter range of ∼7 ± 1.6 nm for reliable sensing (**Figure 3b**). After establishing this optimal range, we evaluated whether the target amplicons produced distinguishable signals. However, as shown in Supplementary **Figure S1**, significant signal overlap was observed between the two targets using conventional statistical analysis. This limitation prompted the adoption of a DNN-based approach for accurate event classification.

To enhance ML performance, we focused on accurate feature extraction and careful DNN architecture design. Among nine tested ML models, the DNN model outperformed others due to the complexity of our signal data. As shown in **Figure 4**, our DNN architecture consists of five hidden layers with 512, 256, 128, 64, and 32 nodes, respectively, each using ReLU activation to mitigate vanishing gradient issues during backpropagation. We trained the model using the AdamW optimizer on 60% of the 2,450 total data points, with 20% used for validation and the remaining 20% reserved for testing. The model achieved 94.2% accuracy on test data, corresponding to a 5.8% error rate at the single-molecule level, demonstrating its reliability in classifying unknown RPA signals.

Further, we validated the performance of the DNN model using a separate set of 35 previously unseen samples, which were completely excluded from the training, validation, and testing phases. This evaluation included independent T_1_, T_2_, and T_1_ + T_2_ samples to assess the model’s true positive classification ability under real-world conditions. As shown in **Figure 5**, the model achieved 100% classification accuracy, confirming its effectiveness at the population level. ROC analysis further demonstrated perfect discriminatory power, with an area under the curve (AUC) of 1.0 for both T_1_, T_2_ classifiactions. Importantly, the event caller performance remained consistent across nanopores of varying diameters (5.45 nm, 6.34 nm, and 7.25 nm), emphasizing the robustness of the approach. These findings establish the utility of this DNN-based classification framework for reliable and scalable molecular diagnostics and suggest its strong potential for detecting co-infections or performing multiplexed target identification in clinical settings.

## CONCLUSIONS

This work presents a robust, probe-free, and deep learning–aided detection platform that significantly advances the field of multiplexed diagnostics. By integrating a novel length-encoded RPA assay with solid-state nanopore sensing and deep neural network (DNN)-based signal classification, we demonstrate accurate, rapid, and scalable detection of both single and multiplex targets without the need for complex optical systems or costly fluorescent probes. The nanopore diameter (∼7 nm) enabled reliable discrimination of RPA amplicons across a broad size range (80– 500 bp), addressing a key limitation in nanopore-based resolution. The DNN classifier, trained on event-level features, achieved 94.2% accuracy in single-molecule classification and 100% accuracy in population-level blind sample identification, with perfect AUC-ROC performance. This current duplex strategy is expandable to multiplex detection, enhancing analytical performance and offering strong potential for miniaturized diagnostics in low-resource settings.

## MATERIALS AND METHODS

### Materials and chemicals

Synthetic Mpox DNA (Quantitative Synthetic Monkeypox virus DNA, VR-3270SD™, 100 μL, 2.8 × 10⁵ copies/μL) was obtained from ATCC Bio, while Human RNase P DNA (Hs_RPP30 Positive Control, #10006626) was sourced from IDT. DNA purification was carried out using a Qiagen kit (#28104). Key materials, including the 1 kb DNA Ladder (#N3232S), Gel Loading Dye, Purple (6X) (#B7025), and nuclease-free water (B1500), were purchased from New England Biolabs Inc. (NEB). Additional reagents, such as SYBR™ Safe DNA Gel Stain, Ultra Low Range DNA Ladder (#10597012), TaqMan™ Fast Virus 1-Step Multiplex Master Mix (#5555532), and Tris-EDTA buffer, were acquired from Thermo Fisher and Sigma-Aldrich. RPA reaction kits and enzymes (freeze-dried pellet tubes; #TALQBAS01) were provided by TwistDx Ltd (Cambridge, UK). Experimental components, including quartz capillaries (0.5 mm inner diameter and 1 mm outer diameter; Sutter Instrument, USA), silver wire (0.008 inches, 25 feet), and accessories like pipette holders (#663040), seals (#647110), and pin connectors (#647110), were sourced from A-M Systems. A microinjector equipped with 34-gauge needles was also purchased from World Precision Instruments.

### RPA assay

#### Design

For the synthetic DNA sequences, we utilized the F3L gene of the monkeypox virus, spanning nucleotides 46,337 to 46,453 (ID: MPXV_USA_2022_OR0002, GenBank accession: ON563414.3), along with the RNase P target sequence sourced from Integrated DNA Technologies (IDT), as detailed in Supplementary **Section 6**. Primers were specifically designed to align with these target regions, with sequence specificity validated through a BLAST search (https://blast.ncbi.nlm.nih.gov/Blast.cgi). Primer quality was further evaluated using the PrimerQuest Tool of IDT and TwistDx design protocols to ensure the absence of primer-dimer formation and secondary structures such as hairpins. All primers, listed in Supplementary **Table S1**, were commercially synthesized and purified.

#### RPA singleplex assay

The RPA reaction, with a total volume of 25 µL, incorporates 1.2 µL of each forward (F_1_ or F_2_) and reverse primers (R_1_ or R_2_) with10 µM initial concentrations, 12.5 µL of 2x reaction buffer, 2.5 µL of 10x basic E-mix, 2 µL of 10 mM dNTPs, 1.25 µL of 20x core reaction mixture, 2.1 µL of nuclease-free water, 1 µL of target sample (Either Mpox or RNase P). We have used the same concentration (10^4^ copies/µL) of both targets. Lastly, 1.25 µL of MgAcO (280 mM) was used to prevent premature reaction initiation. The master mix is incubated at 39°C for 20 minutes. The detailed recipe is provided in Supplementary **Table S2**.

#### RPA multiplex assay

The RPA reaction, with a total volume of 25 µL, incorporates 1.2 µL of each forward (F_1_ and F_2_) and reverse primer (R_1_ and R_2_) with 500 nM and 400 nM final concentrations.^49^ Other reagents such as 12.5 µL of 2x reaction buffer, 2.5 µL of 10x basic E-mix, 1 µL of 400 µM final concentrations of dNTPs, and 1.25 µL of 20x core reaction mixture. 1 µL of each target sample (10^4^ copies/µL Mpox and 10^4^ copies/µL RNase P). Lastly, 1.25 µL of MgAcO (280 mM) was used to prevent premature reaction initiation. The master mix is incubated at 39°C for 20 minutes. To terminate the reaction, heat it to 85°C for 1 min and cool it to 4°C.^50,51^ The detailed recipe is provided in Supplementary **Table S3**.

#### Gel analysis of RPA products

Post-RPA, the amplicons are extracted from the 5% agarose gel electrophoresis performed at 8 V/cm using SYBR™ Safe DNA Gel Stain. Residual RPA components were removed using the QIAquick Gel Extraction Kit. The required bands are then analyzed using the BIO-RAD GelDoc Go Imaging System with a 20-second exposure.

### q-PCR assay

In this study, a one-step RT-qPCR approach was employed to screen RNase P primers for use in a duplex assay. Each 20 µL reaction mixture contained 10 µL of TaqMan Fast Virus 1-Step Master Mix, 1.2 µL each of forward and reverse primers (final concentration: 0.6 µM), 0.5 µL of probe (final concentration: 0.25 µM), 1 µL of RNase P template, and 6.1 µL of PCR-grade water. The RT-qPCR was carried out under standard thermal cycling conditions, starting with an initial denaturation at 95 °C for 20 seconds, followed by 40 amplification cycles. Each cycle included a denaturation step at 95 °C for 3 seconds and an annealing/extension step at 60 °C for 30 seconds. The detailed recipe is provided in Supplementary **Table S4**.

### Nanopore fabrication, data acquisition, and analysis

#### Nanopore fabrication

Quartz capillaries were cleaned in Piranha solution (H₂SO₄:H₂O₂, 3:1) at 85°C for 30 minutes, then rinsed with deionized water and dried at 120°C for 20 minutes in a vacuum oven. Nanopores were fabricated using a laser pipette puller (P-2000, Sutter Instruments, USA) under a single-stage protocol (heat 650, filament 4, velocity 60, delay 145, pull 160–215).^52^ The finished pores, typically 3.3-12.4 nm in diameter, were filled with either 1 M KCl or 2 M LiCl in Tris-EDTA buffer (pH 8.0) and filtered through a 0.2 µm syringe filter (WHA67802502, Whatman).^41^ Pore characterization was carried out by performing current-voltage (I–V) sweeps from −800 to +800 mV.^52,53^

#### Nanopore measurements

Following methods similar to our previous work,^7,39^ a stable voltage was applied using a 6363 DAQ card (National Instruments, USA), and the ionic current was amplified with an Axopatch 200B amplifier (Molecular Devices, USA). Data were sampled at 100 kHz and low-pass filtered at 10 kHz through LabVIEW. To reduce external noise, all measurements were conducted inside a custom Faraday cage mounted on an optical table (Thorlabs). Two Ag/AgCl electrodes connected to the amplifier were placed in the cis and trans chambers (with the cis side grounded), each containing the salt solution. Amplicons were diluted 20-fold before measurement (Supplementary **Figure S7**) to prevent clogging; if clogging did occur, five cycles of voltage sweeps from −800 to +800 mV were performed to clear the pore.

#### Data analysis

Ionic current traces were evaluated for signal-to-noise ratios (SNR) above 3, dwell times under 10 ms, and interarrival times of 200 ms. Ionic current traces were acquired at two sampling frequencies based on experimental needs: 50 kHz for high-resolution *I–t* trace and 1.67 kHz for long-*I–t* trace. For individual event shape visualization and data storage, recordings were resampled to 1 kHz as needed. Translocation events were characterized by extracting dwell times, current blockades, contour plots, and histograms using Python 3.8.3 and MATLAB R2020b. Data visualization was carried out with OriginPro 2021 and Python. For size-based counting, a constant 400 mV was applied to the nanopore via the 6363 DAQ card.

#### DNN model and event caller

Data for the DNN model was collected by performing nanopore measurements on each sample for 2–5 minutes, depending on pore clogging, with a goal of capturing at least 100-300 events per sample.^54^ Precisely controlling a 7 nm pore size is challenging for reproducibility, so we used nanopores within the ∼7 ± 1.6 nm range for data collection to train the DNN model. Each recorded current trace was then analyzed to identify translocation events and extract features such as dwell time and current drop. These events were labeled according to the known sample type (e.g., either Mpox or control), forming a labeled dataset. In cases of severe clogging, fewer events were recorded. Measurements were conducted using 15 nanopores for training (Supplementary **Figure S10**) and testing datasets, with an additional 15 nanopores used to collect completely unseen data for model application. All datasets, including training, testing, and untrained unknown samples, were saved and are provided in the supplementary materials.

## ASSOCIATED CONTENT

The Supporting Information is available and it includes supplemental figures, tables, and codes.

## DATA AVAILABILITY

Data from the study is available from the corresponding author upon reasonable request.

## ACKNOWLEDGMENTS

This work was partially supported by the National Institutes of Health (R33AI147419) and the National Science Foundation (2045169, 2319913). Any opinions, findings, conclusions, or recommendations expressed in this work are those of the authors and do not necessarily reflect the views of the National Science Foundation and National Institutes of Health.

## DECLARATIONS

The authors declare no competing financial interest.

## AUTHOR INFORMATION

### Corrosponding Author

Weihua Guan-Department of Intelligent Systems Engineering, Indiana University, Bloomington, IN, 47408, United States;

### Authors

Md. Ahasan Ahamed-Department of Intelligent Systems Engineering, Indiana University, Bloomington, IN, 47408, United States; and Department of Electrical Engineering, Pennsylvania State University, University Park, PA, 16802, United States;

Muhammad Asad Ullah Khalid-Department of Intelligent Systems Engineering, Indiana University, Bloomington, IN, 47408, United States;

Anthony J. Politza-Department of Biomedical Engineering, Pennsylvania State University, University Park, PA, 16802, United States;

Wahid Uz Zaman-of Computer Science and Engineering, Pennsylvania State University, University Park, PA, 16802, United States;

### Author Contributions

M.A.A. designed and validated the assay, conducted investigations, acquired data, performed formal analysis, conducted data analysis, and wrote the manuscript. M.A.K. handled data visualization, acquired data, and performed data analysis. A.P.J. wrote the necessary programming code and contributed to the review and editing. W.U.Z. gave the initial idea to work with DNN and provide insights into model optimization. W.G. developed the concept, provided resources, and supervised the overall experimental design.

## TOC FIGURE

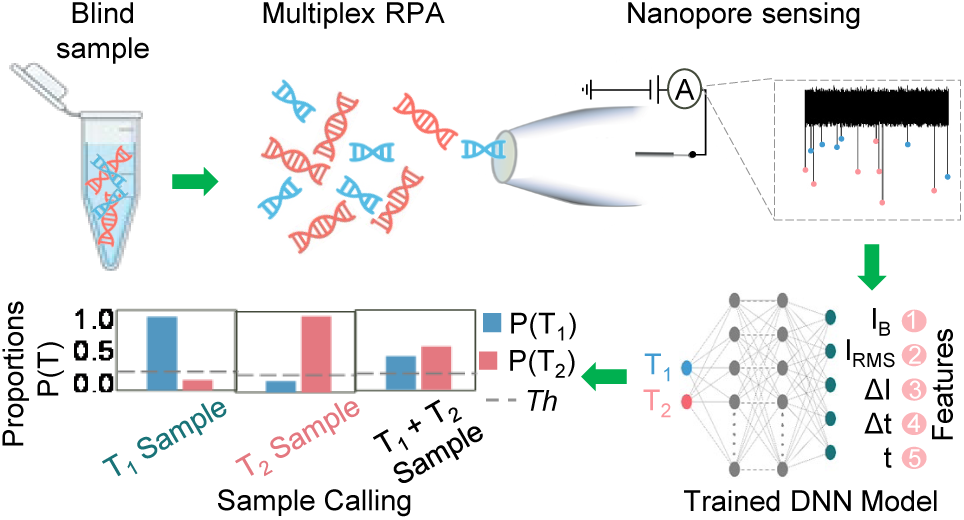
Multiplex RPA amplicons are classified via nanopore sensing and deep learning, enabling rapid, probe-free molecular diagnostics at the single-molecule level.

